# Cancer cells suppress NK activity by actin-driven polarisation of inhibitory ligands at the synapse

**DOI:** 10.1101/2025.02.03.636217

**Authors:** Céline Hoffmann, Liza Filali, Hannah Wurzer, Diogo Pereira Fernandes, Takouhie Mgrditchian, Flora Moreau, Max Krecké, Clément Thomas

## Abstract

Natural killer (NK) cells engage target cells via the immunological synapse, where inhibitory and activating signals determine whether NK cell cytotoxicity is suppressed or activated. We report that cancer cells can rapidly remodel their actin cytoskeleton upon NK cell engagement, leading to F-actin accumulation at the synapse. This process inhibits NK cell activation as indicated by impaired MTOC and lytic granule polarization. Exploring the underlying mechanism, we found that actin remodelling drives the recruitment of inhibitory ligands, such as HLA-A, -B, and -C, to the synapse. Disrupting HLA interaction with their cognate inhibitory receptors KIRs restored NK cell activation. Using NK cells expressing inhibitory KIR receptors, matched or unmatched to HLA molecules on cancer cells, we show that synaptic F-actin accumulation and matching KIR-HLA interactions jointly suppress NK cell cytotoxicity. Our findings reveal a novel immune evasion strategy in which cancer cells impair NK cell activation by altering synaptic signalling through actin cytoskeleton-driven recruitment of inhibitory signals to the immunological synapse.

## Introduction

Natural killer (NK) cells are cytotoxic lymphocytes that play a crucial role in cancer immunosurveillance. Unlike CD8+ T cells, which require prior sensitization to specific antigens and clonal expansion, NK cells can recognize and eliminate abnormal cells upon first encounter, positioning them a critical first line of defence against tumorigenesis. Instead of antigen-specific receptors, NK cells express a repertoire of germline-encoded inhibitory and activating receptors that collectively guide their response (1). Activating receptors detect stress-induced or abnormal ligands on target cells, pushing NK cells toward cytotoxic activation, while inhibitory receptors recognize self-molecules signalling cellular normalcy and restraining NK cell activation (1,2). The decision of NK cells to kill or spare potential targets is ultimately determined by the balance of inhibitory and activating signals received through these receptors. This finely tuned interplay enables NK cells to mount a cytotoxic response against the diverse phenotypes exhibited by cancer cells while preventing inappropriate cytotoxic responses against healthy cells. Although NK cells are primarily associated with the innate immune system, they can also display adaptive properties which may be harnessed for cancer immunotherapy (1,3).

Healthy cells typically express high levels of major histocompatibility complex class I (MHC I) molecules on their surface, which serve as “self” markers. These molecules play a critical role in promoting NK cell tolerance by engaging the main inhibitory receptors, including inhibitory killer-cell immunoglobulin-like receptors (iKIRs) and the CD94/NKG2A heterodimeric receptor (4–6). In contrast, tumours frequently downregulate MHC-I molecules on their surface, reducing detection by CD8+ T cells, which require MHC-I to present cancer cell antigens for immune recognition. This reduction in self signals (or “missing-self”) weakens the inhibitory input mediated by iKIRs and NKG2A, making NK cells more prone to activation (7). However, NK cell activation also requires additional inputs from activating receptors, such as NKG2D and the natural cytotoxicity receptors (NCRs), including NKp30, NKp44 and NKp46 (8). For instance, stress molecules like MICA, MICB and UL16-binding proteins (ULBPs) are frequently overexpressed on tumour cells, serving as potent activating ligands for NKG2D. Additionally, the stress-induced self-molecule B7-H6 is upregulated in various cancers and specifically engages NKp30, leading to robust NK cell activation (9). When an activating receptor binds to its ligand, intracellular signalling is initiated through distinct motifs in adaptor molecules (6). Immunoreceptor tyrosine-based activation motif (ITAM)-bearing adaptors, such as DAP12, rely on phosphorylation by Src family kinases to create docking sites for downstream signalling kinases, such as Syk and Zap70. In contrast, ITAM-independent activating receptors like NKG2D signal through adaptors, such as DAP10 which contains a YINM motif (10). Phosphorylation of this motif recruits PI3K and other signalling proteins. While both DAP12 and DAP10 use different signalling intermediates, both pathways converge on actin cytoskeleton polymerization and reorganization (11,12). Conversely, engagement of inhibitory receptors with their ligands leads to phosphorylation of immunoreceptor tyrosine-based inhibition motifs (ITIMs) within their cytoplasmic domain by Src family kinases (6). Phosphorylated ITIMs recruit of phosphatases, particularly Src homology 2-domain-containing protein tyrosine phosphatase-1 (SHP-1) and SHP-2, which dephosphorylate key molecules in the activation pathway, thereby suppressing the activation cascade (6,13,14). In addition, an increasing number of studies support that NK cells are also regulated by non-MHC-I specific immune checkpoints, such as PD-1, LAG-3, TIM-3 and TIGIT, which have been reported to induce NK cell functional exhaustion (15–18). However, the consistency and prevalence of these pathways in directly modulating NK cell anti-tumour activity across various human cancers are still under evaluation.

The recognition of cancer cells and their subsequent killing by NK cells rely on the formation a stable cell-to-cell contact known as the lytic immunological synapse (IS). This process initiates with NK cell adhesion to the target cell, typically mediated by the interaction between lymphocyte function-associated antigen-1 (LFA-1) on NK cells and intracellular adhesion molecules (ICAMs) on the cancer cell surface (19). This adhesion is strengthened by the engagement of activating receptors, leading to the reorganization of the actin cytoskeleton into a circular network of filaments at the periphery of the IS, and the polarization of the microtubule organizing centre (MTOC) and associated cytotoxic granules toward the target cell (19–22). In addition to releasing classical cytotoxic granules containing perforins, granzymes and other cytotoxic molecules, NK cells have recently been shown to secrete membraneless particles termed supramolecular attack particles (SMAPs) containing perforin and granzyme B surrounded by a thrombospondin-1 shell (23). The actin cytoskeleton plays multiple and critical roles in NK cell-mediated cytotoxicity, supporting IS initiation, stabilization and maturation into distinct functional domains (24,25). During degranulation, actin dynamics regulate the formation of localized clearances within the synaptic cortical F-actin meshwork, allowing the granules to pass through and fuse with the cell membrane (26–30). Additionally, actin dynamics contribute to the regulation of synaptic signalling through various mechanisms, including the generation of forces that modulate receptor-ligand interactions, the regulation of the conformation and activity of critical checkpoint molecules such as SHP-1, and the assembly and transport of signalling clusters to specific regions of the IS (12,31,32).

While the actin cytoskeleton is widely recognized as a critical and multifunctional component of ISs formed by NK cells and CD8+ T cells with their targets, its role(s) within the attacked cancer cells remain comparatively understudied (24). Nevertheless, growing evidence indicates that the configuration and dynamic remodelling of the actin cytoskeleton in cancer cells during interactions with cytotoxic lymphocytes critically influence IS outcomes (24,33). The cortical actin cytoskeleton of cancer cells—a key regulator of their mechanical properties—plays a pivotal role in generating synaptic forces that modulates the activation of mechanosensitive immunoreceptors and the pore-forming activity of perforin (34–38). In addition, our previous work established that accumulation of F-actin at the cancer cell side of the IS during interactions with NK cells triggers resistance to NK cell-mediated cytotoxicity (39,40). This synaptic evasion strategy is conserved across multiple breast cancer and chronic lymphocytic leukaemia (CLL) cell lines and has been validated in primary CLL patient samples. However, the molecular mechanism by which actin cytoskeleton remodelling protects cancer cells during NK cell attack remains unknown.

## Results

### Synaptic polarization of the cancer cell actin cytoskeleton inhibits MTOC and cytotoxic granule polarization in primary NK cells

We previously established that the rapid and sustained polarization of F-actin at the immunological synapse (IS) in cancer cells during interactions with NK cells significantly enhances their resistance to NK cell-mediated cytotoxicity (39,40). Notably, cancer cells undergoing actin cytoskeleton remodelling exhibited reduced levels of NK cell-delivered granzyme B, suggesting a potential disruption in NK cell cytotoxic activation. To further investigate this hypothesis, we assessed NK cell activation status during interactions with MDA-MB-231 breast cancer cells, comparing targets with or without F-actin synaptic polarization. Primary NK cells were isolated from healthy donors and used as effector cells. Activation status was evaluated by visualizing the microtubule-organizing centre (MTOC) via γ-tubulin staining and quantifying its proximity to the IS by measuring the distance to its centre (Figure 1A and C). Additionally, cytotoxic granules were detected using granzyme B staining, and their polarization was assessed as the percentage of granzyme B localized within the third of the NK cell closest to the IS, or “synaptic area” (Figure 1B and C). For each donor-derived NK cell preparation, 40 cell-cell conjugates were analysed via confocal microscopy (20 for each target cell actin phenotype). Our data reveal a statistically significant and striking difference in NK cell polarization depending on the F-actin configuration in target cells. Notably, NK cells interacting with target cells exhibiting synaptic F-actin accumulation showed a dramatic reduction in the polarization of their MTOC and granules compared to those engaging with target cells lacking actin cytoskeleton remodelling (Figure 1C-E). To assess whether the NK cell polarization defect is dependent on iKIR signalling, we extended our analysis using NK-92MI cells, an IL-2-independent variant of the NK-92 cell line that lacks iKIRs and is therefore unresponsiveness to MHC-I (41). Unlike primary NK cells, these effector cells displayed a similarly pronounced polarization of both their MTOC and granules toward target cells, irrespective of the target cell’s actin phenotype (n=60; p=ns; Figure 1F-H). Taken together, these findings underscore the pivotal role of actin cytoskeleton remodelling in determining whether MDA-MB-231 cells adopt an inhibitory or stimulatory phenotype toward primary NK cells and suggest that this outcome depends on inhibitory signalling, particularly involving iKIRs.

**Figure 1:**
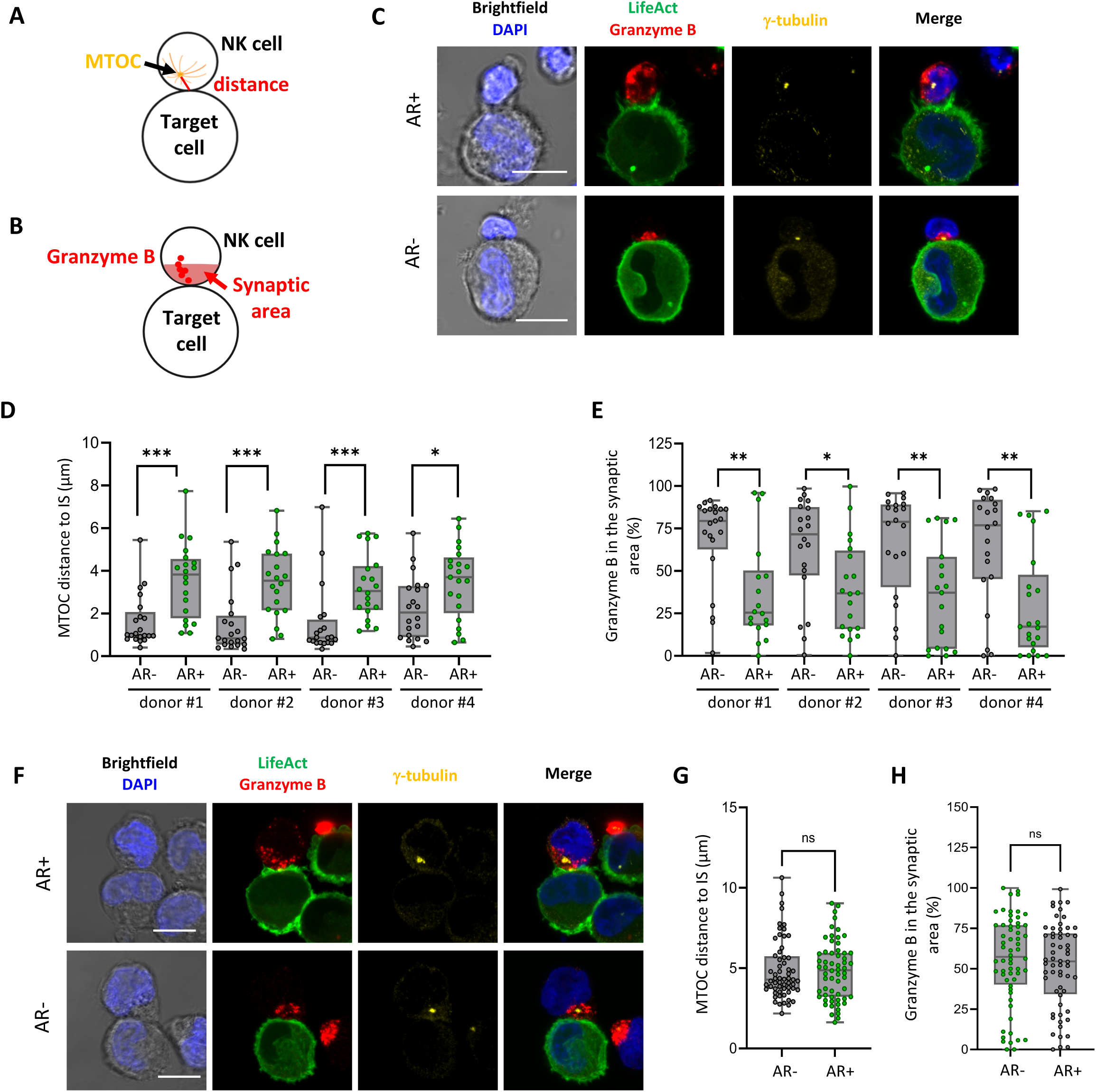
F-actin polarization at the cancer cell side of the immunological synapse is associated with impaired lytic machinery polarization in primary NK cells. Emerald-LifeAct-expressing MDA-MB-231 cells (green) were co-cultured with NK cells for 60 minutes, followed by immunolabeling for Granzyme B, γ-Tubulin along with DAPI staining for the nucleus. **(A-B)** NK cell lytic machinery polarization was assessed by measuring the distance between the MTOC and the IS centre **(A)**, and by quantifying the proportion of granzyme B localized within the synaptic area, defined as the proximal third of the cell closest to the IS **(B). (C)** Representative confocal images (maximum intensity projection of 4 optical slices) of cell-to-cell conjugates between primary NK cells and MDA-MB-231 cells, with or without synaptic actin cytoskeleton remodelling (AR+ and AR-, respectively). **(D-E)** Quantitative analysis of primary NK cell lytic machinery polarization, collected from NK cells isolated from 4 distinct donors, with n=20 cell-to-cell conjugates analysed per condition. Data on the MTOC distance from the IS **(D)** and Granzyme B enrichment at the IS **(E)** are presented. **(F)** Representative confocal images of cell-to-cell conjugates between NK92-MI cells and AR+ or AR-MDA-MB-231 cells. **(G-H)** Quantitative analysis of NK92-MI cell lytic machinery polarization, collected from 3 independent experiments, with n=60 cell-to-cell conjugates analysed per condition. Data on the MTOC distance from the IS **(G)** and granzyme B enrichment at the IS **(H)** are presented. Statistical significance was determined using the Mann-Whitney test. Scale bars: 10 µm.

### Synaptic polarization of the cancer cell actin cytoskeleton correlates with localized enrichment of surface HLA inhibitory ligands

Building on the above findings, we investigated the distribution of classical MHC-I molecules (HLA-A, -B, -C), the main inhibitory ligands for NK cells, on the surface of target cells conjugated with primary NK cells. High-resolution confocal microscopy revealed a marked accumulation of HLA-A, -B, - C molecules at the IS in target cells exhibiting synaptic actin cytoskeleton polarization (Figure 2A). This synaptic enrichment resulted in a several-fold increase in fluorescence intensity at the IS compared to the non-synaptic regions of the same cell. Conversely, target cells lacking synaptic F-actin polarization displayed a relatively uniform distribution of HLA-A, -B, -C molecules, with occasional reductions in signal intensity at the IS. To increase the statistical power of the analysis, imaging flow cytometry was employed to examine a large cohort of NK cell-target cell conjugates using primary NK cells from two independent donors. Specific synaptic and non-synaptic masks were applied to quantify the synaptic enrichment of F-actin and HLA-A, -B, -C molecules in target cells, as described previously (42) (Supplementary Figure 1). Target cells with a polarized actin cytoskeleton showed a significantly higher frequency and magnitude of HLA-A, -B, -C enrichment at the IS compared to those lacking F-actin polarization (p < 0.001, n = 100 target cells per phenotype; Figure 2B and C). This relationship was further supported by strong correlation values between synaptic F-actin polarization and HLA-A, -B, - C accumulation (R= 0.7 and R = 0.8 for NK cells from donor #4 and #5 respectively, p < 0.001; Figure 2D). Interestingly, the overall surface levels of HLA-A, -B, -C molecules remained comparable between target cells with and without synaptic actin polarization (Figure 2E), suggesting that synaptic enrichment results from the redistribution of pre-existing surface molecules. To assess whether this redistribution depends on interactions with cognate NK cell receptors, we repeated the analysis using iKIR-deficient NK-92MI cells as effector cells. Both confocal microscopy and imaging flow cytometry yielded similar results to those obtained with primary NK cells (Supplementary Figure 2A-E). Specifically, HLA-A, -B, -C molecules accumulated prominently at the IS in target cells exhibiting synaptic F-actin polarization, whereas their distribution remained uniform in target cells lacking this phenotype (Supplementary Figure 2A-C). Notably, a strong correlation was also observed between synaptic F-actin polarization and HLA-A, -B, -C accumulation in assays with NK-92MI, reaching 0.86 (p < 0.001; Supplementary Figure 2D).

**Figure 2:**
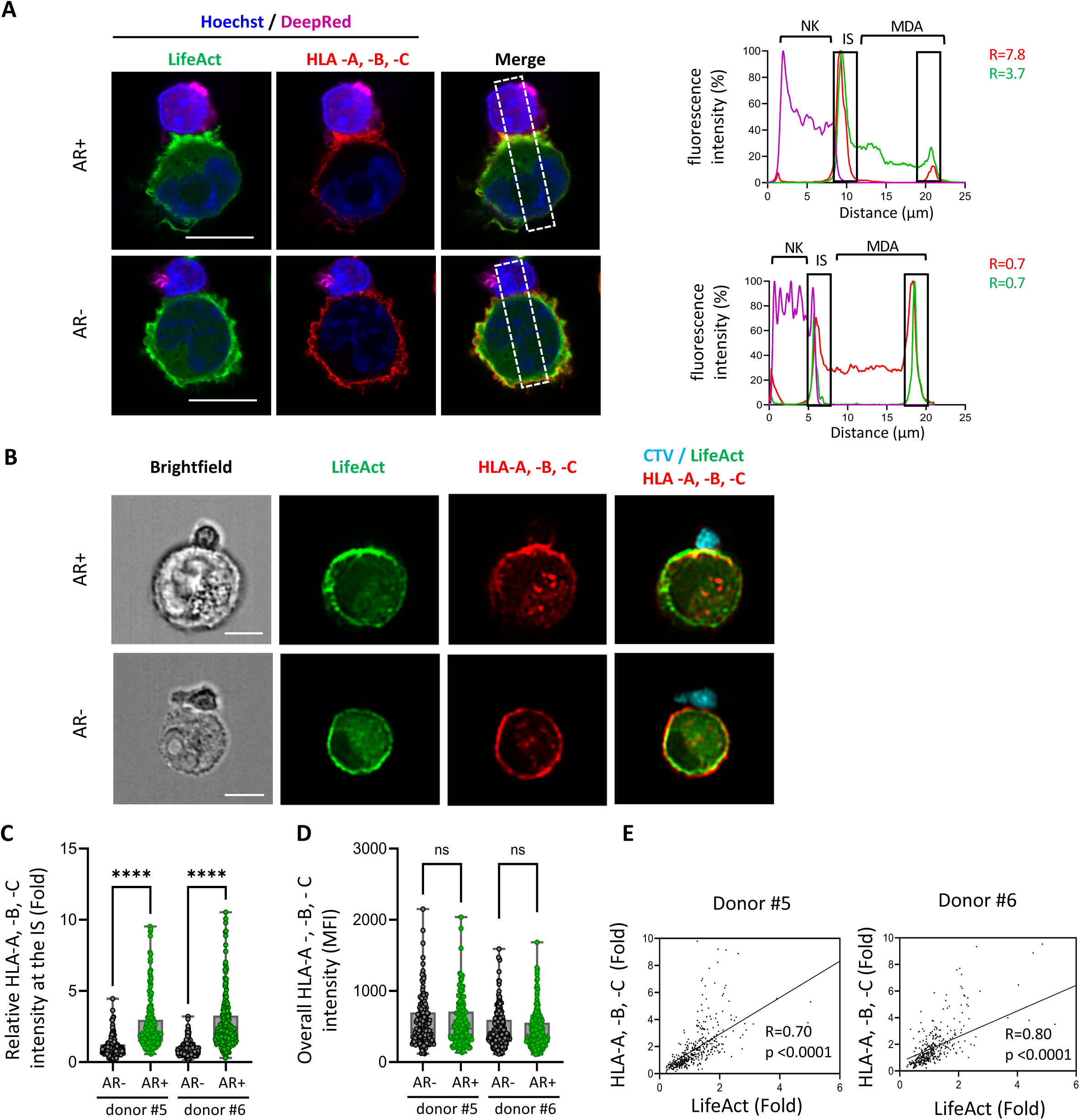
Polarization of the actin cytoskeleton at the cancer cell side of the immunological synapse correlates with the local accumulation of HLA molecules during interaction with NK cells. **(A)** Emerald-LifeAct-expressing MDA-MB-231 cells (green) were pre-labelled for HLA-A, -B, -C (red) and co-cultured for 40 minutes with Deep-Red-stained primary NK cells (purple), along with Hoechst staining for the nucleus (blue). Representative Airyscan images show cell-to-cell conjugates formed between primary NK cells and MDA-MB-231 cells with or without synaptic actin cytoskeleton remodelling (AR+ and AR-, respectively) . The dashed rectangle indicates the 50-pixel-wide line used to measure mean fluorescence intensity (MFI) for Emerald-LifeAct, HLA-A, -B, -C and Deep Red signals. The upper chart shows MFI profiles for AR+ MDA-MB-231 cells, while the lower chart displays profiles for AR-MDA-MB-231 cells. The relative intensity of HLA-A, -B, -C (red), Emerald-LifeAct (green) and Deep Red (purple) at the IS is displayed on the plot. **(B-E)** Emerald-LifeAct-expressing MDA-MB-231 cells (green) were pre-labelled for HLA-A, -B, -C (red) and co-cultured for 40 minutes with CTV-stained primary NK cells (cyan). Data were collected from NK cells isolated from 2 distinct donors, with n=200 cell-to-cell conjugates analysed per condition. **(B)** Representative imaging flow cytometry images of cell-to-cell conjugates between primary NK cells and AR+ or AR-MDA-MB-231 cells. **(C)** Emerald-LifeAct relative intensity at the IS was used to classify MDA-MB-231 cells into AR+ and AR-groups (ratio >1 and ratio <1 respectively). Relative HLA-A, -B, -C intensities at the IS in these 2 subgroups are presented. **(D)** The overall MFI of HLA-A, -B, -C across the entire cell membrane of the cancer cell was measured in cell-to-cell conjugates formed between primary NK cells and AR+ or AR-MDA-MB-231 cells. Statistical significance was determined using the Mann-Whitney test. **(E)** Correlation graph showing the relative intensities of Emerald-LifeAct and HLA-A, -B, -C at the IS across the entire population of cell-to-cell conjugates analysed in (C and D), without distinguishing between AR+ or AR-MDA-MB-231 cells. The correlation was determined using Spearman’s correlation coefficient. Scale bars: 10 µm.

To determine whether our findings extend beyond MDA-MB-231 cells, we examined the synaptic abundance of HLA-A, -B, -C molecules in MDA-MB-468 cells, a more epithelial-like and EGFR-dependent breast carcinoma cell line. When MDA-MB-468 cells were engaged by either primary NK cells or NK-92MI cells, synaptic F-actin polarization similarly promoted HLA-A, -B, -C accumulation at the IS, confirming that this phenomenon is not limited to MDA-MB-231 cells (Supplementary Figure 3A, B, D and E). Moreover, in these cells as well, overall HLA-A, -B, -C surface expression remained unchanged regardless of synaptic actin polarization (Supplementary Figure 3C and F), further supporting the idea that IS enrichment results from redistribution rather than increased expression.

Together, these findings indicate a strong association between synaptic polarization of the cancer cell actin cytoskeleton and the localized enrichment of HLA-A, -B, -C molecules at the IS. This enrichment appears to occur independently of inhibitory KIR interactions, as demonstrated by its persistence in assays with KIR-deficient NK-92MI cells, and is driven by the polarization of pre-existing surface molecules toward the IS.

### Polarization of HLA inhibitory ligands to the immunological synapse requires a functional actin cytoskeleton

The role of the actin cytoskeleton in the polarization of HLA-A, -B, -C molecules to the IS was investigated using mycalolide B (MycB), a marine macrolide toxin that depolymerizes F-actin into monomeric actin and irreversibly disrupts actin dynamics (43). Treatment with 1 µM MycB resulted in extensive F-actin depolymerization in MDA-MB-231 cells (Supplementary Figure 4A). Following MycB pre-treatment and subsequent drug removal, MDA-MB-231 cells were co-cultured with NK cells as previously. Quantitative analysis using imaging flow cytometry revealed a marked reduction in the percentage of MycB-treated cells exhibiting actin cytoskeleton polarization during NK cell interactions, decreasing from approximately 45% in DMSO-treated control cells to less than 10% (p < 0.001; n = X) (Figure 3A). This disruption of synaptic actin remodelling led to a statistically significant and substantial decrease in the synaptic enrichment of HLA-A, -B, -C molecules compared to control cells (p < 0.0001; Figure 3B). To verify that these effects were specific to actin cytoskeleton disruption, additional experiments were conducted using colchicine, a microtubule-disrupting agent. Treatment of MDA-MB-231 with 2.5 µM colchicine effectively disrupted microtubules (Supplementary Figure 4B) but had no measurable impact on the percentage of cells displaying synaptic actin polarization during NK cell interactions, which remained around 45% (Figure 3C). Furthermore, unlike MycB, colchicine did not significantly impair the enrichment of HLA-A, -B, -C molecules at the IS (Figure 3D). These findings underscore the critical and specific role of the actin cytoskeleton in mediating the recruitment of HLA-A, -B, -C molecules to the IS.

**Figure 3:**
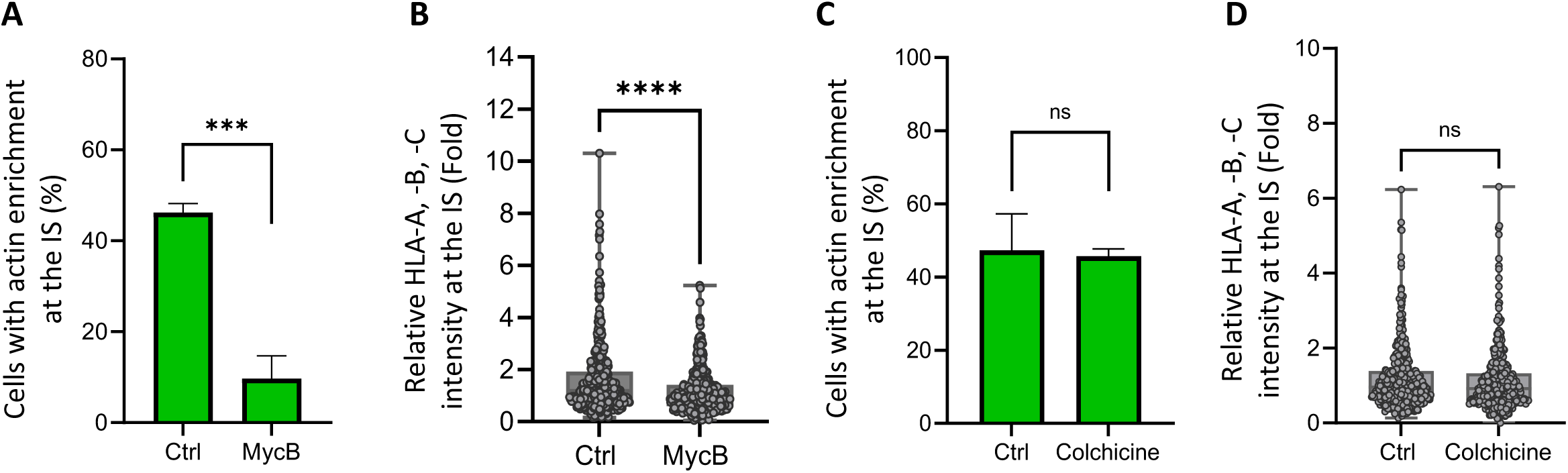
Synaptic polarization of HLA-A, -B, -C depends on the actin cytoskeleton, not microtubules. Emerald-LifeAct-expressing MDA-MB-231 cells (green) were treated with cytoskeletal-disrupting agents and labelled for HLA-A, -B, -C (red), conjugated with CTV-stained NK92-MI cells (cyan) for 40 minutes, and analysed by imaging flow cytometry. **(A-B)** Cancer cells were treated with Mycalolide B (an inhibitor of actin filament polymerization) at a concentration of 1 µM for 20 minutes. **(A)** Relative Emerald-LifeAct intensity at the immunological synapse was quantified, and the percentage of cell-to-cell conjugates with synaptic actin cytoskeleton remodelling is shown. **(B)** Relative HLA-A, -B, -C intensities at the IS in cell-to-cell conjugates are presented. **(C-D)** Cancer cells were treated with colchicine (a microtubule-disrupting agent) at a concentration of 2.5 µM for 90 minutes. **(C)** Relative Emerald-LifeAct intensity at the immunological synapse was quantified, and the percentage of cell-to-cell conjugates with synaptic actin cytoskeleton remodelling is shown. **(D)** Relative HLA-A, -B, -C intensities at the immunological synapse in cell-to-cell conjugates are presented. Data were collected from three independent experiments, with n=200 cell-to-cell conjugates analysed per condition. Statistical significance was determined using the Mann-Whitney test.

### Actin cytoskeleton remodelling-driven synaptic polarization of inhibitory ligands enhances inhibitory signalling toward NK cells

Based on our findings, we hypothesized that inhibition of NK cell polarization in front of target cells exhibiting synaptic F-actin polarization results from the recruitment and local accumulation of HLA inhibitory ligands, which in turn enhances inhibitory signalling towards NK cells. To test this hypothesis, we investigated the interplay between synaptic actin cytoskeleton remodelling and HLA inhibitory ligand polarization in mediating NK cell suppression, employing an HLA-blocking antibody to disrupt iKIR signalling. The HLA-blocking antibody was first validated to ensure that the localization pattern of HLA molecules, including their synaptic accumulation in cancer cells with F-actin synaptic polarization, remained consistent with the observations made using non-blocking antibodies (Supplementary Figure 5).

MDA-MB-231 cells were pre-incubated with the HLA-blocking antibody or an IgG isotype control before co-culture with primary NK cells. Polarization of the MTOC and cytotoxic granules in NK cells derived from two different donors was quantified using imaging flow cytometry (Figure 4). HLA blockade, but not the IgG control, significantly increased the polarization of both the MTOC and cytotoxic granules in NK cells interacting with target cells exhibiting synaptic actin cytoskeleton remodelling. Notably, under these conditions, NK cell polarization was restored to levels comparable to those observed in interactions with target cells lacking actin cytoskeleton remodelling (Figure 4B and C). We extended our analysis to MDA-MB-468 cancer cells, where HLA blockade similarly reversed actin cytoskeleton-driven inhibition of NK cell polarization (Supplementary Figure 6A and B). These findings reinforce and expand upon previous observations, supporting that the protective effect of F-actin synaptic polarization in cancer cells against NK cell cytotoxicity is mediated by inhibitory signalling through iKIR engagement with HLA molecules.

**Figure 4:**
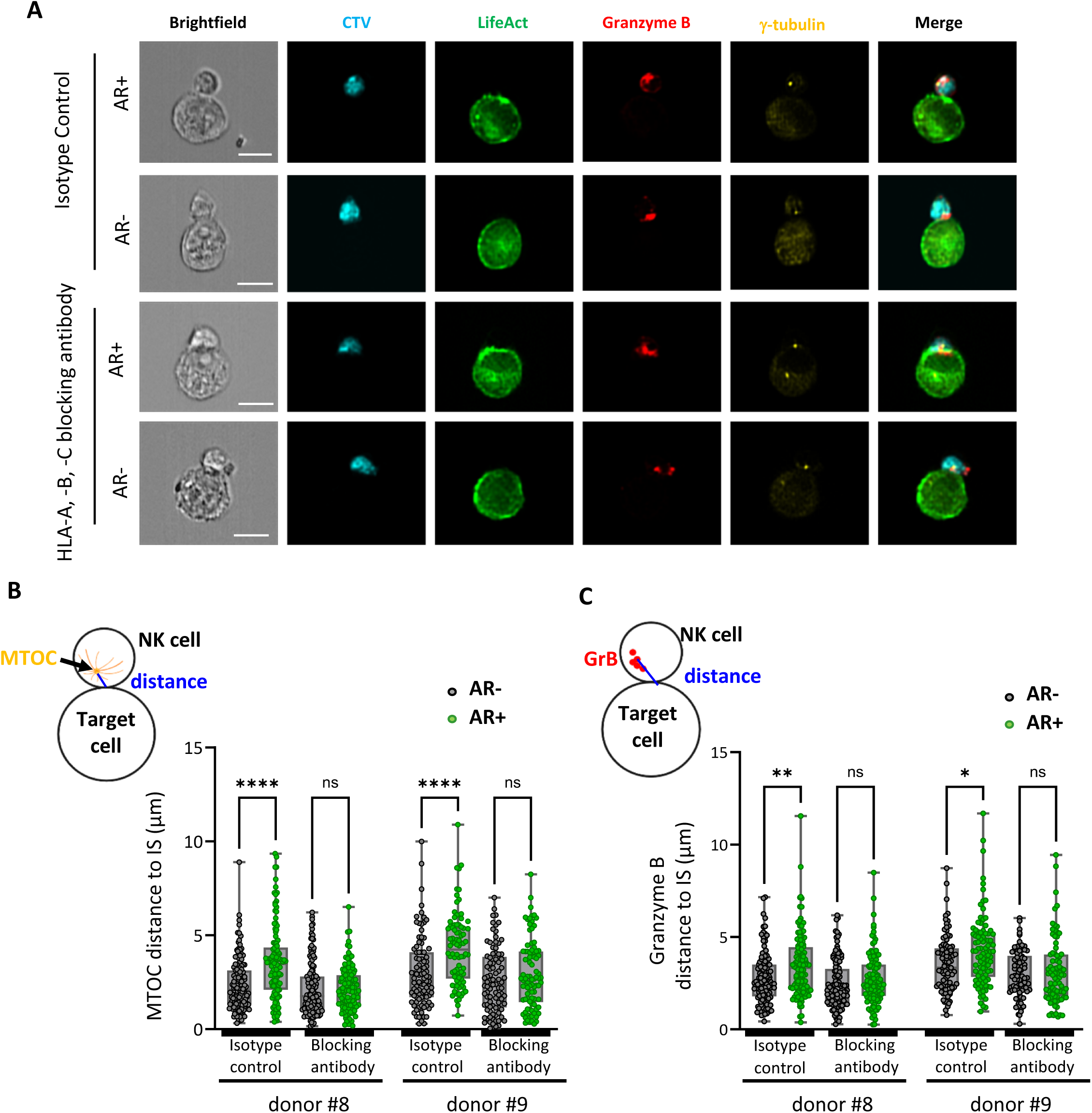
HLA-blocking antibody counteracts synaptic actin remodelling-driven inhibition of NK cell polarization. Emerald-LifeAct-expressing MDA-MB-231 cells (green) were pre-treated with either a HLA-A, -B, -C blocking antibody or an isotype control antibody before being conjugated with CTV-labelled primary NK cells (cyan) for 60 minutes. After conjugation, cells were immunolabelled for Granzyme B and γ-Tubulin and analysed using imaging flow cytometry. Data were collected from NK cells isolated from 2 distinct donors, with n=100 cell-to-cell conjugates analysed per condition. **(A)** Representative imaging flow cytometry images of cell-to-cell conjugates between primary NK cells and MDA-MB-231 cells, with or without synaptic actin cytoskeleton remodelling (AR+ and AR-, respectively). Scale bar: 10 µm. **(B-C)** Emerald-LifeAct relative intensity at the immunological synapse was used to classify MDA-MB-231 cells into AR+ and AR-groups (ratio >1 and ratio <1 respectively). **(B)** NK cell lytic machinery polarization was evaluated by measuring the distance between the MTOC and the synapse center. **(C)** The distance between the granzyme B centroid and the synapse centre was measured. Distances for AR+ and AR-subgroups are presented. Statistical significance was determined using the Mann-Whitney test.

To further determine whether actin cytoskeleton remodelling in cancer cells impairs NK cell activation through inhibitory ligand-receptor interactions, we employed matched and mismatched iKIR-ligand models. NK cells from donors were isolated and sorted into subpopulations either expressing or lacking KIR2DL1 (Supplementary Figure 7A), an iKIR that recognizes group 2 HLA-C (HLA-C2) molecules expressed by MDA-MB-231 cells (44). Both subpopulations (KIR2DL1+ and KIR2DL1-) were expanded to ensure sufficient cell numbers for subsequent analyses. The expression of HLA-C2 on the surface of MDA-MB-231 cell was confirmed using a specific antibody (Supplementary Figure 7B). Notably, HLA-C2 expression was found to be relatively low compared to the overall expression of HLA-A,-B,-C (Supplementary Figure 7B and C). HLA-C was confirmed to polarize to the synaptic region in NK cell-conjugated target cells exhibiting synaptic F-actin accumulation, whereas it remained uniformly distributed on the surface of target cells lacking actin cytoskeleton remodelling (Supplementary Figure 7D).

Imaging flow cytometry was used to quantitatively assess MTOC and cytotoxic granule polarization in KIR2DL1-and KIR2DL1+ NK cells from three different donors. NK cells from each subpopulation were analysed during interactions with target cells either exhibiting or lacking actin cytoskeleton remodelling (Figure 5). KIR2DL1-NK cells demonstrated comparable MTOC and cytotoxic granule polarization regardless of the target cell’s cytoskeletal phenotype. In contrast, KIR2DL1+ NK cells exhibited polarization similar to that of KIR2DL1-NK cells when interacting with target cells lacking actin cytoskeleton remodelling but showed significantly reduced polarization when engaging target cells with synaptic F-actin accumulation. These findings indicate that actin-driven polarization of inhibitory ligands to the cancer cell side of the IS strengthens inhibitory signalling toward interacting NK cells, leading to suppression of NK cell polarization.

**Figure 5:**
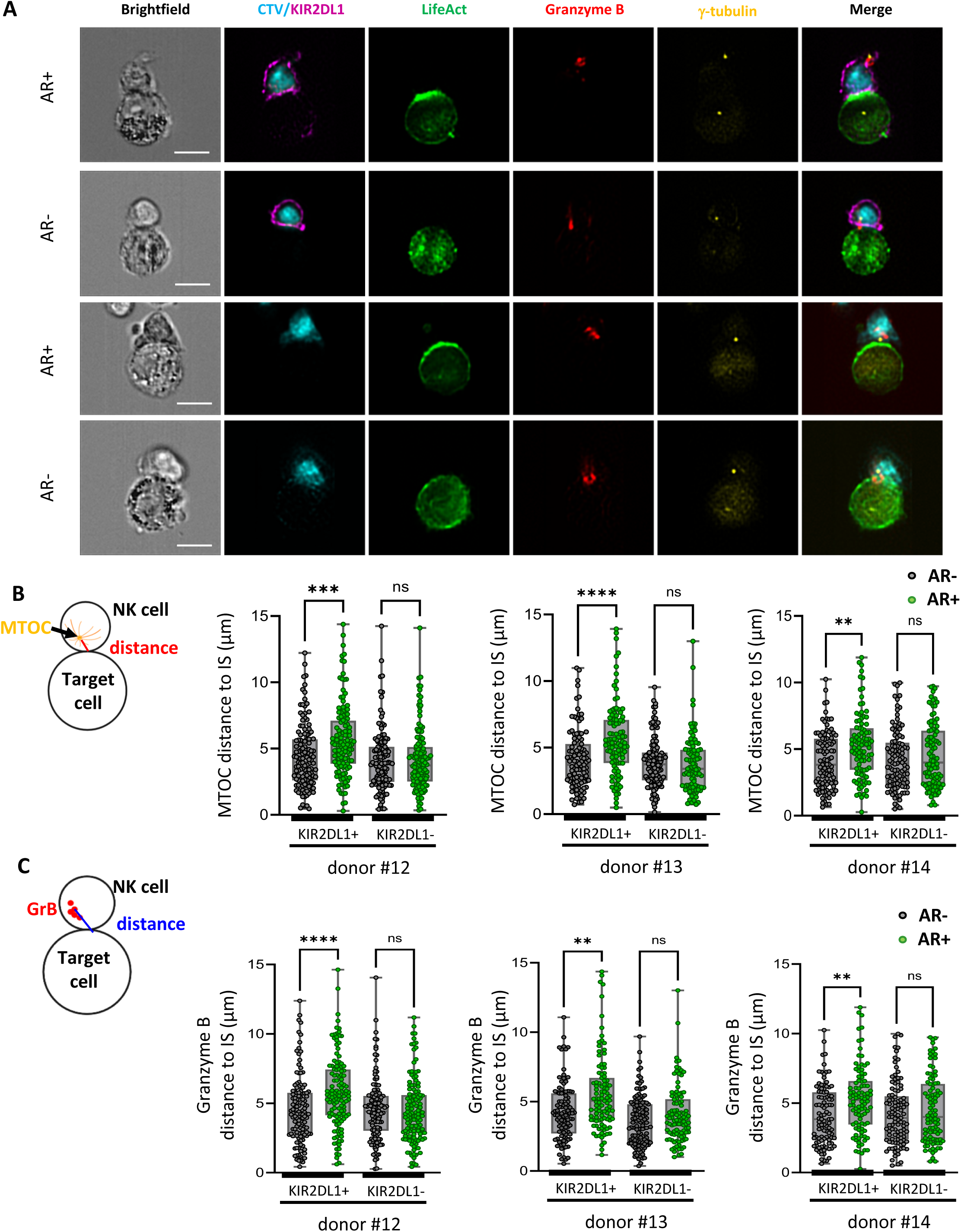
Inhibition of NK cell polarization by synaptic actin polarization in cancer cells is driven by cognate HLA-iKIR interactions. Emerald-LifeAct-expressing MDA-MB-231 cells (green) were co-cultured for 60 minutes with CTV-labelled primary NK cells, either KIR2DL1^+^ or KIR2DL1^-^ cells. After co-culture, cells were immunolabelled for granzyme B (red), γ-Tubulin (yellow) and KIR2DL1 (purple). Data were collected from NK cells isolated from 3 distinct donors, with n=100 cell-to-cell conjugates analysed per condition. **(A)** Representative imaging flow cytometry images of cell-to-cell conjugates between primary NK cells (KIR2DL1^+^ or KIR2DL1^-^) and MDA-MB-231 cells, with or without synaptic actin cytoskeleton remodelling (AR+ and AR-, respectively). Scale bar: 10 µm. **(B-C)** Emerald-LifeAct relative intensity at the synapse was used to classify MDA-MB-231 cells into AR+ and AR-groups (ratio >1 and ratio <1, respectively). **(B)** NK cell lytic machinery polarization was evaluated by measuring the distance between the MTOC and the synapse centre. **(C)** The distance between the granzyme B centroid and the IS centre was measured. Distances for AR+ and AR-subgroups are presented. Statistical significance was determined using the Mann-Whitney test.

## Discussion

MHC-I downregulation is a well-established mechanism by which tumour cells evade cytotoxic T cell responses and resist immunotherapy (45). Paradoxically, this reduction in MHC-I expression increases their susceptibility to NK cell attack, as MHC-I molecules play a crucial role in suppressing NK cell activation (7). Our findings reveal that cancer cells can counteract this vulnerability by redistributing their limited MHC-I pool to the IS through actin remodelling. By locally concentrating MHC-I at the IS, cancer cells amplify inhibitory signalling to interacting NK cells, effectively mimicking higher MHC-I expression and thereby dampening NK cell activation. Notably, a similar mechanism has been described in mature dendritic cells (DCs), which engage NK cells to stimulate them (46) while avoiding destruction (47). In this so-called regulatory synapse (48), F-actin polymerization at the DC interface facilitates MHC-I accumulation, reinforcing inhibitory signalling and suppressing NK cell cytotoxicity. Our findings suggest that cancer cells exploit this protective mechanism, originally evolved to modulate immune responses, as a mean to evade NK cell-mediated killing. We demonstrate that even a relatively low abundance of MHC-I molecules, such as HLA-C2 on MDA-MB-231 cells, can amplify inhibitory signalling in a measurable manner when strategically concentrated at the synapse— precisely at the optimal location and timing during NK cell interactions.

Recent studies have identified the target cell’s actin cytoskeleton as a key regulator of cytotoxic lymphocyte activity by modulating target cell rigidity and deformability (34,38). These biophysical properties contribute to the mechanical forces at the IS, which influence force-sensitive receptors on cytotoxic lymphocytes, fine-tuning immune responses through a process known as mechanosurveillance (38). Our findings extend this understanding by demonstrating that, beyond shaping the biophysical properties of target cells, the actin cytoskeleton directly regulates the molecular composition of the IS by controlling the synaptic density and distribution of key surface molecules. Interestingly, the synaptic polarization of HLA molecules is strictly actin-dependent and independent of microtubules. In addition, this process occurs irrespective of the engagement of HLA molecules with their cognate receptors on NK cells, as evidenced by the persistence of F-actin polarization even when HLA-iKIR interactions are pharmacologically inhibited or when NK92-MI cells, which lack iKIRs, are used. Therefore, actin dynamics emerge as the primary driver in redirecting inhibitory signalling toward the IS. While directly inhibiting actin dynamics is not a viable therapeutic approach due to its high toxicity and widespread cellular effects, targeting the molecular trigger on NK cells that drives rapid F-actin remodelling in target cells could offer new opportunities for enhancing NK cell-mediated anti-tumour responses. Additionally, neutralizing this trigger through CAR-NK cell engineering could improve therapy efficacy by offering a more specific alternative to broadly overriding inhibitory signalling, which carries a higher risk of off-target toxicity (49). Identifying the molecular trigger responsible for actin remodelling is therefore a critical next step towards the translation of our findings into therapeutic strategies.

At the same time, dissecting the functional consequences of this remodelling is essential to fully understand how it shapes immune interactions. A key question in this regard is how synaptic actin remodelling mediates local increase in MHC-I surface density. At the DC-NK cell regulatory synapse, F-actin has been proposed to stabilize the low-affinity interaction between MHC-I and KIRs (47). By analogy to its role in regulating MHC-II diffusion (50), the actin-based membrane skeleton may create membrane compartments or ’fences’ that constrain molecular mobility, thereby increasing MHC-I concentration at the synapse (47). Since our data indicate that the overall MHC-I surface expression does not significantly differ between target cells with or without polarization, we also favour a model in which the actin cytoskeleton regulates the distribution of pre-existing MHCI molecules, e.g. by restricting their diffusion away from the synaptic region, rather than facilitating their targeted delivery to the synapse.

Beyond clustering inhibitory ligands, emerging evidence suggests that actin remodelling at the cancer cell side of the IS plays a broader role in reconfiguring intracellular trafficking. Specifically, actin dynamics have been recently implicated in the enrichment and fusion of exosome-containing multivesicular bodies at the IS (51), suggesting that this interface functions as a polarized secretory domain. While the role of targeted secretion from the cancer cell side of the IS remains to be fully elucidated, it may contribute to immune evasion by directing exosomes enriched in MHC-I and immune checkpoint molecules toward NK cells, thereby reinforcing inhibitory signalling and sustaining NK cell suppression. Other recent studies have further emphasized the dynamic nature of the cancer cell side of the IS. In melanoma cells, a calcium-dependent membrane repair response is rapidly activated following sub-lethal hits by CD8+ T cells, involving lysosome mobilization to the IS to counteract cytotoxicity (52). In breast cancer cells under NK cell attack, the synaptic membrane composition undergoes remodelling, characterized by an increase in densely packed lipids, which reduces perforin binding and enhances resistance to NK cell killing (53). Given the well-established roles of actin in regulating membrane trafficking, it is worth investigating whether synaptic actin remodelling directly orchestrates these defence mechanisms, potentially serving as a point of convergence that could be targeted to simultaneously inhibit multiple evasion strategies.

Collectively, our findings reveal a deeper level of organization at the IS, suggesting that in its resistant state, the cancer cell side of the IS mirrors the lymphocyte’s side, with each cell actively polarizing its molecular arsenal for either attack or defence. Just as cytotoxic lymphocytes undergo actin-driven polarization to coordinate effector functions, cancer cells seem to exploit cytoskeletal remodelling to reinforce inhibitory signalling, repair and modify the membrane, and potentially direct immunosuppressive secretions. These insights emphasize the IS as a dynamic battlefield, where the structural and molecular adaptations of each interacting cell ultimately dictate the outcome.

## Methods

### Cell Lines

MDA-MB-231 (ATCC HTB-26), MDA-MB-468 (ATCC HTB-132), and NK-92MI (ATCC CRL-2408) cell lines were purchased from the American Type Culture Collection (ATCC). Breast cancer cell lines were transduced to stably express the mEmerald-Lifeact-7 F-actin reporter (Addgene, plasmid #54148), as previously described (39). MDA-MB-231 and MDA-MB-468 cells were maintained in Dulbecco’s Modified Eagle Medium (DMEM) supplemented with 10% foetal bovine serum (FBS), 100 U/mL penicillin, and 100 μg/mL streptomycin. NK-92MI cells were grown in RPMI-1640 medium supplemented with 10% FBS, 10% horse serum, 100 U/mL penicillin, and 100 μg/mL streptomycin. All cell lines were authenticated and verified to be free of cross-contamination via short tandem repeat (STR) profiling (Microsynth). Cells were cultured in a humidified incubator at 37°C with 5% CO₂ and were routinely screened for Mycoplasma contamination.

### Isolation of human primary NK cells and amplification of the KIR2DL1^+^/^-^ subpopulations

Primary NK cells were isolated from cryopreserved peripheral blood mononuclear cells (PBMCs) obtained from buffy coats of healthy, anonymous donors, provided by the Luxembourg Red Cross. NK cell isolation was performed using the human NK cell isolation kit (Miltenyi Biotec #130-092-657) via negative selection, according to the manufacturer’s instructions. Briefly, PBMCs were incubated with biotin-conjugated monoclonal antibodies against antigens not expressed by NK cells, followed by a second incubation with microbeads conjugated to monoclonal antibodies. Labelled cells were subsequently separated using a magnetic column system. After isolation, the NK cells were washed and resuspended in pre-warmed RPMI medium supplemented with 10% FBS, 1% HEPES (10 mM), 1% MEM-NEAA, 1% sodium pyruvate, 1% penicillin/streptomycin, IL-2 (100 IU/mL, Miltenyi Biotec #130-097-746) and IL-15 (10 ng/mL, Miltenyi Biotec #130-095-765). The cells were adjusted to a final concentration of 2 x 10⁶ cells/mL and incubated overnight before being used in experiments. In a subset of experiments, isolated human primary NK cells were stained to identify, isolate and amplify KIR2DL1-expressing cells. Live NK cells were identified by incubating them with the Zombie NIR fixable viability kit (BioLegend #423106) for 15 minutes at room temperature. After washing in MACS buffer (Miltenyi Biotec #130-091-221), Fc receptor blocking solution (Human TruStain FcX, BioLegend #422302) was added for 10 minutes to minimize nonspecific binding. Cells were then washed and incubated for 30 minutes in the following antibody staining mix: FITC-conjugated anti-CD56 antibody (BioLegend #318304), PB-conjugated anti-CD3 antibody (BioLegend #300330), and PE-conjugated anti-CD158a (KIR2DL1) antibody (Miltenyi #130-120-446). KIR2DL1^+^ and KIR2DL1^-^ NK cells were sorted using a fluorescence-activated cell sorter (BD FACSymphony^TM^ S6 lite). Sorted NK cells were seeded in 96-well U-bottom plates at a density of 8 x 10^4^ cells/well in 200 µL of complete medium containing 60% DMEM, 25% F-12, 10% human serum, 1% HEPES (10 mM), 1% MEM-NEAA, 1% penicillin/streptomycin, IL-2 (400 IU/mL, Miltenyi Biotec #130-097-746) and PHA-P (1 µg/mL, Invivogen #inh-phap). To facilitate cell expansion, the NK cells were co-cultured with 1 x 10^5^ of PBMCs per well from 2 distinct donors. To block PBMCs cell proliferation, they were pre-treated with mitomycin C (20 µg/mL, Merck #M4287-2MG) during 30 minutes at 37°C in complete media prior to co-culture with NK cells. Cells were split every 3 days over a two-week period before use in downstream experiments.

### Fluorescence staining and preparation of cell-to-cell conjugates for confocal microscopy

NK cells were stained with DeepRed dyes (Invitrogen #C34565, 1:1000 dilution) in serum-free medium for 20 minutes at 37°C. Ligand labeling was performed in MACS buffer (Miltenyi Biotec #130-091-221) using PE-conjugated anti-HLA-A, -B, -C (BD Pharmingen #560964) or PE-conjugated anti-HLA-C (BD Pharmingen #566372) antibodies for 30 minutes at 4°C. For conjugate formation, equal numbers of labelled NK cells (7 × 10⁴ in 100 µL) and cancer cells (7 × 10⁴ in 100 µL) were mixed and incubated at 37°C for 40 minutes (for HLA localization) or 60 minutes (for NK activation) in the presence of Hoechst 33342 (1:2000, Miltenyi Biotec #130-111-569). Conjugates were then transferred onto poly-L-lysine-coated µ-slide chambers (Ibidi #80806) and allowed to adhere for 8 minutes. Cells were fixed with 4% paraformaldehyde (PFA, Agar Scientific) for 15 minutes at room temperature and permeabilized with 0.1% Triton X-100 for 5 minutes. Following a 1-hour blocking step in PBS containing 5% BSA and 10% FBS, cells were incubated with primary antibodies against Granzyme B (BioLegend #396406) and γ-Tubulin (Santa Cruz Biotechnology #sc-17788) for 2 hours at room temperature. After washing, secondary antibodies (Alexa Fluor-conjugated goat anti-mouse, anti-rabbit, or anti-rat, e.g., AF555 or AF647, Invitrogen) were applied. Finally, cells were washed, and the medium was replaced with mounting medium (Ibidi #50001) for imaging. In some experiments, adherent target cells were pretreated with Mycalolide B (1 µM, Enzo #BML-T123-0020) for 20 minutes, colchicine (2.5 µM, Merck #C9754) for 1.5 hours, or DMSO as a control. After fixation and permeabilization, cells were stained with an AF647-conjugated anti-α-Tubulin antibody (BioLegend #627908) to assess drug effects.

### Confocal imaging and quantitative image analysis

Images were acquired using a Zeiss LSM880 fast Airyscan confocal microscope with excitation lasers (405 nm, 488 nm, 543 nm, 594 nm, 633 nm) in a multitrack setup. Acquisitions were performed in confocal or Airyscan mode, with z-stacks (0.2 μm interval) followed by deconvolution using Zen Blue v3.5. Image type (confocal/Airyscan) is specified in the figure legends. For Granzyme B analysis, a z-stack of the entire NK cell was acquired (0.5 μm intervals, ∼20 slices). Quantification was performed using ImageJ v1.53t. Granzyme B polarization was assessed using a Z-projection (‘sum slices’ method) as previously described (51). A region of interest (ROI) encompassing the target cell was manually segmented, with the synaptic region defined as the one-third portion in direct contact with the NK cell. Granzyme B localization was quantified as the ratio of mean pixel intensity (MPI) in the synaptic ROI to the total ROI. To measure MTOC positioning relative to the synapse, coordinates of the MTOC and synapse centre were obtained using the ‘multipoint tool,’ and distances were calculated as:

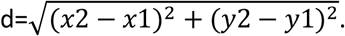

For Emerald LifeAct and HLA fluorescence profiling, a 100-pixel (3.5 µm) line was drawn across conjugates, and fluorescence intensity was analysed using the ‘Plot Profile’ tool. Enrichment was determined as the ratio of maximum intensity at the IS to the maximum intensity at the opposite side of the target cell.

### Fluorescence staining and preparation of cell-to-cell conjugates for imaging flow cytometry (IFC)

NK cells were stained with CellTrace Violet (CTV, Thermo Fisher #C34557, 1:8000) and Zombie NiR (BioLegend #423105, 1:2000) for 20 minutes in PBS before conjugation. For extracellular labeling, cells were incubated in MACS buffer (Miltenyi Biotec #130-092-987) with PE/Dazzle594-conjugated anti-HLA-A, -B, -C (BioLegend #311410) or PE-conjugated anti-CD158a (BioLegend #374904) for 30 minutes at 4°C. Target and effector cells were conjugated at a 2:1 ratio (6 × 10⁵ NK cells and 3 × 10⁵ MDA-MB-231 cells in 200 µL). After 40 minutes (for HLA localization) or 60 minutes (for NK activation), cells were fixed with 4% PFA, permeabilized with 0.1% Triton X-100, and stained with AF594-conjugated anti-γ-Tubulin (Santa Cruz Biotechnology #sc-17788) and APC-conjugated anti-Granzyme B (BioLegend #396408). Cells were resuspended in 35 µL PBS for acquisition. For HLA-A, -B, -C blockade, MDA-MB-231 cells were pre-incubated with Ultra-LEAF™ anti-HLA-A, -B, -C (BioLegend #311428) or an isotype control (BioLegend #400264) at 50 µg/mL for 30 minutes, followed by co-culture with primary NK cells. Fixation and intracellular staining for Granzyme B and MTOC followed. To verify antibody fixation, one sample was stained with goat anti-mouse AF555. For cytoskeleton studies, target cells were detached and treated with Mycalolide B (1 µM, 20 min), colchicine (2.5 µM, 1.5 h), or DMSO. Post-treatment, cells were washed and stained for HLA-A, -B, -C. Conjugation time was reduced to 30 minutes for colchicine-treated samples due to the reversible nature of colchicine’s effects.

### Imaging flow cytometry and quantitative image analysis

Samples were acquired using an ImageStream®X Mark II (Cytek Biosciences) with four lasers (405 nm, 488 nm, 561 nm, 642 nm) and operated using INSPIRE® software. Single-color controls were used for matrix compensation. A total of 5 × 10⁴ to 8 × 10⁴ conjugates were acquired per sample at 60X magnification. IFC analysis was conducted using IDEAS software, following Biolato et al. (42) Briefly, the synaptic mask was generated at the NK-target interface and extended into the target cell (∼1/3 of total target area). A non-synaptic mask covering the remaining portion of the cell was also created. MPI was calculated within each mask, and F-actin enrichment was determined as the ratio of MPI in the synaptic mask to the non-synaptic mask. Cells were categorized as AR− (ratio <1) or AR+ (ratio >1), with classification validated through visual inspection. The same analysis was applied to ligand enrichment, restricting masks to the cell membrane. MTOC and Granzyme B polarization were assessed using the ’delta centroid’ feature, calculating the distance between the centroid of the MTOC or Granzyme B mask and the synaptic mask centroid.

### HLA expression analysis by flow cytometry

MDA-MB-231 cells were labelled with the Zombie NIR viability kit (BioLegend #423106, 15 min, RT). After washing, cells were incubated with PE-conjugated anti-HLA-A, -B, -C (BD Pharmingen #560964), anti-HLA-C (BD Pharmingen #566372), or their respective isotype controls (PE Mouse IgG1, ĸ Isotype control; BD Pharmingen #554680 or PE Mouse IgG2b, ĸ Isotype control; BD Pharmingen #555058) for 30 minutes at 4°C in the dark. Samples were analysed using a NovoCyte Quanteon 4025 flow cytometer, and HLA expression was assessed using FlowJo software.

### Statistical analysis

Statistical analyses were performed using GraphPad Prism v10.3.1. Normality was assessed using the Shapiro-Wilk test. Non-parametric tests (Mann-Whitney) were used when data deviated from normality. Proportional comparisons were assessed via a Z-score test for two population proportions (https://www.socscistatistics.com/tests/ztest/). Statistical details are provided in figure legends.

## Acknowledgements.

Clément Thomas’s group is supported by the Luxembourg National Research Fund (FNR), Luxembourg (C21/BM/15752542/SYNAPODIA), the Fondation Cancer, Luxembourg (FC/2019/02/ACTIMMUNE) and Think Pink Luxembourg. Hannah Wurzer is recipient of a PhD fellowship from the Luxembourg National Research Fund (FNR; PRIDE15/10675146/CANBIO). Diogo Pereira Fernandes and Max Krecké are recipients of PhD fellowships from the Fonds De La Recherche Scientifique (FNRS), Belgium (Télévie 7.4594.23 and 7.4531.20, respectively). Takouhie Mgrditchian is recipient of a Postdoctoral fellowship from Fonds De La Recherche Scientifique (FNRS), Belgium (Télévie 7.4597.22). We acknowledge the National Cytometry Platform (NCP) for assistance with the generation of cytometry data. The NCP is supported by funding from Luxembourg’s Ministry of Higher Education and Research (MESR).

## Author contributions

CH Conceptualization, Formal Analysis, Investigation, Methodology, Supervision, Visualization, Writing – original draft. LF Conceptualization, Formal Analysis, Investigation, Methodology, Supervision, Visualization, Writing – original draft. HW Conceptualization, Investigation, Methodology, Writing – review & editing. DPF Investigation, Formal Analysis, Validation. TM Investigation. FM Investigation. MK Investigation, Methodology. CT Conceptualization, Funding acquisition, Project administration, Supervision, Writing – original draft.

---

**Supplementary Figure 1:**
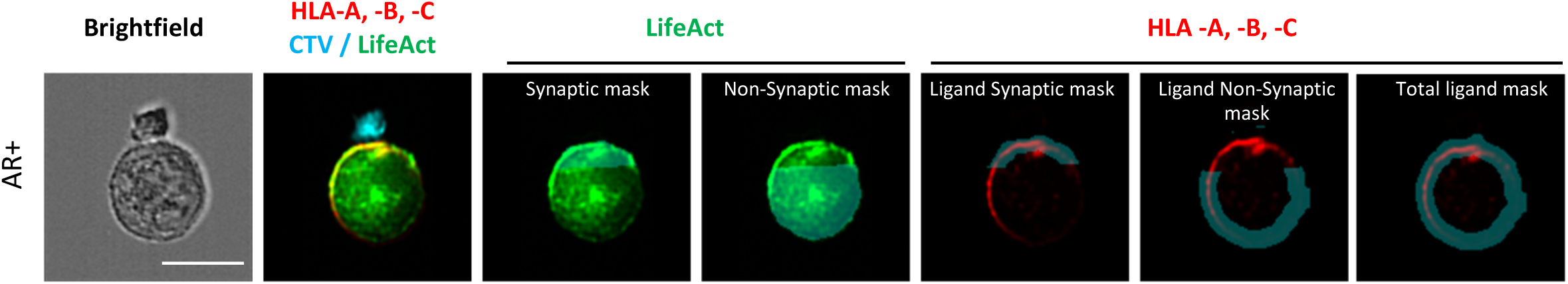
Description of the masks used for imaging flow cytometry analysis. Emerald-LifeAct-expressing MDA-MB-231 cells (green) were pre-labelled for HLA-A, -B, -C (red) and co-cultured for 40 minutes with CTV-stained primary NK cells (cyan). Representative imaging flow cytometry images of cell-to-cell conjugates between primary NK cells and AR+ or AR-MDA-MB-231 cells are presented. A synaptic mask, defined as the proximal third of the cell closest to the synapse, and a non-synaptic mask, encompassing the remaining two-thirds of the cell, were applied. For Emerald-LifeAct intensity measurements, the masks included both the intracellular region and cell membrane. For HLA-A, -B, -C intensity measurements, the masks were restricted to the cell membrane. Relative Emerald-LifeAct and HLA-A, -B, -C, intensities at the IS were calculated as the ratio of mean fluorescence intensity (MFI) in the synaptic mask to the non-synaptic mask. Additionally, a mask covering the entire cell membrane of cell was designed. Masks are shown in light blue. Scale bar: 10 µm.

**Supplementary Figure 2:**
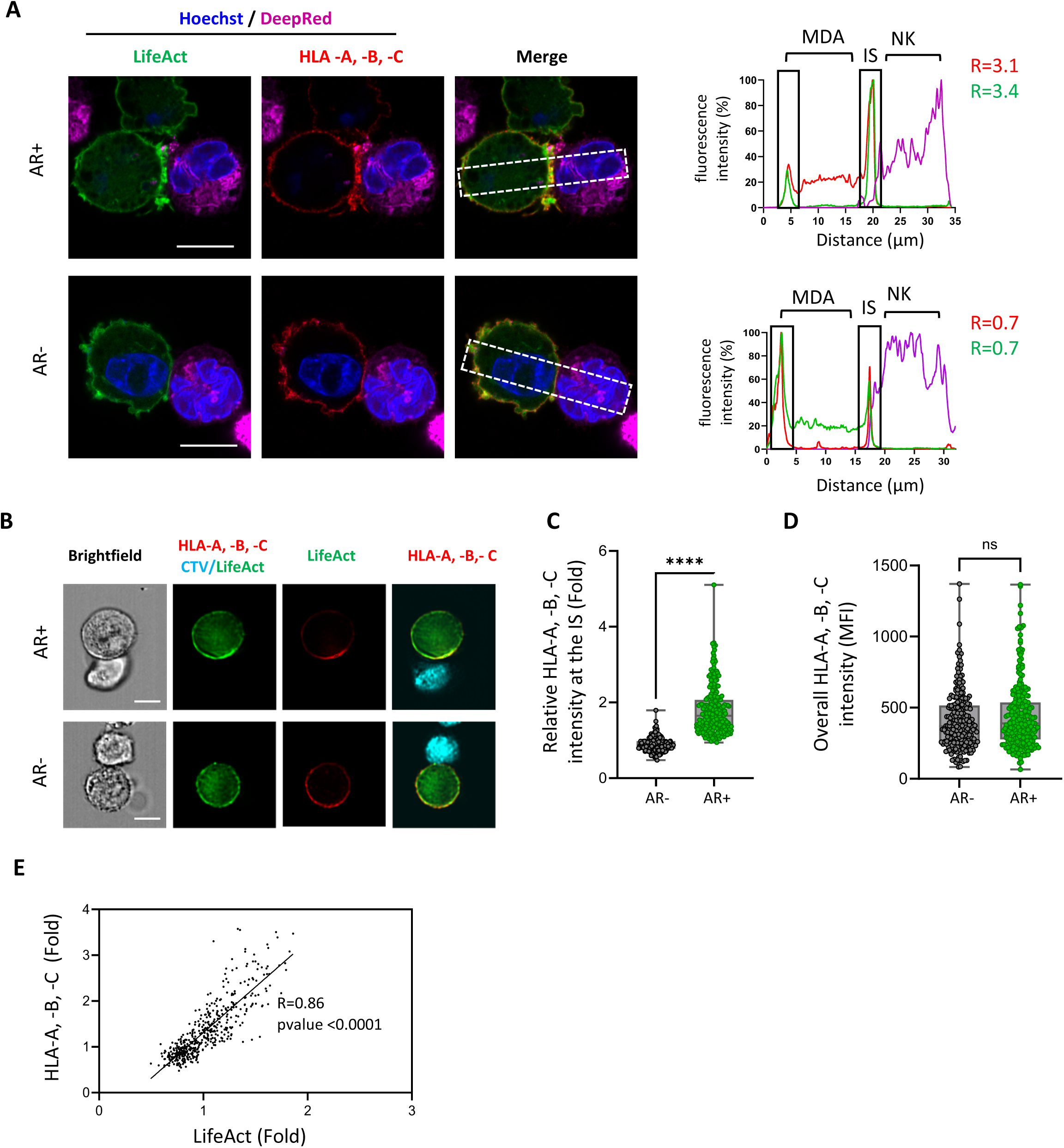
Polarization of the actin cytoskeleton at the cancer cell side of the immunological synapse correlates with the local accumulation of HLA molecules during interaction with NK92-MI cells. **(A)** Emerald-LifeAct-expressing MDA-MB-231 cells (green) were pre-labelled for HLA-A, -B, -C (red) and co-cultured for 40 minutes with Deep-Red-stained NK92-MI cells (purple), along with Hoechst staining for the nucleus (blue). Representative Airyscan images show cell-to-cell conjugates formed between NK92-MI cells and MDA-MB-231 cells with or without synaptic actin cytoskeleton remodelling at the immunological synapse (AR+ and AR-, respectively). The dashed rectangle indicates the 50-pixel-wide line used to measure mean fluorescence intensity (MFI) for Emerald-LifeAct, HLA-A, -B, -C and CMRA signals. The upper chart shows MFI profiles for AR+ MDA-MB-231 cells, while the lower graph displays profiles for AR-MDA-MB-231 cells. The relative intensity of HLA-A, -B, -C (red) and Emerald-LifeAct (green) at the synapse are displayed on the plot. **(B-E)** Emerald-LifeAct-expressing MDA-MB-231 cells (green) were pre-labelled for HLA-A, -B, -C (red) and co-cultured for 40 minutes with CTV-stained NK92-MI cells (cyan). **(B)** Representative imaging flow cytometry images of cell-to-cell conjugates between NK92-MI cells and AR+ or AR-MDA-MB-231 cells. **(C)** Emerald-LifeAct relative intensity at the synapse was used to classify MDA-MB-231 cells into AR+ and AR-groups (ratio >1 and ratio <1 respectively). Relative HLA-A, -B, -C intensities at the IS in these 2 subgroups are presented. **(D)** The overall MFI of HLA-A, -B, -C across the entire cell membrane of the cancer cell was measured in cell-to-cell conjugates formed between NK92-MI and AR+ or AR-MDA-MB-231 cells. Data were collected from 3 independent experiments, with n=250 cell-to-cell conjugates analysed per condition. Statistical significance was determined using the Mann-Whitney test. **(E)** Correlation graph showing the relative Emerald-LifeAct and HLA-A, -B, -C intensities at the IS across the entire population of cell-to-cell conjugates analysed in C-D, without distinguishing between AR+ MDA-MB-231 and AR-MDA-MB-231 cells. The correlation was determined using Spearman’s correlation coefficient. Scale bars: 10 µm.

**Supplementary Figure 3:**
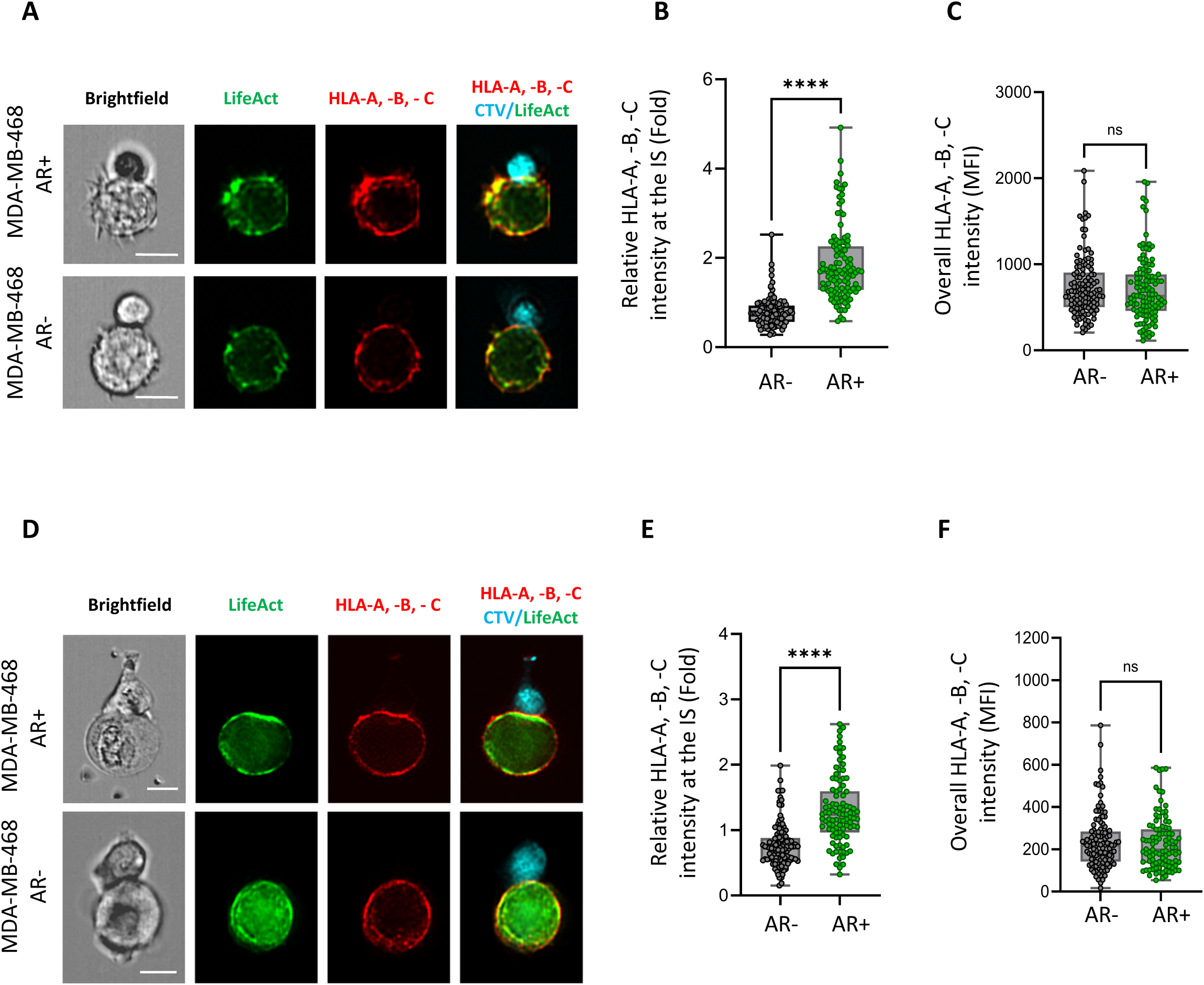
Association between synaptic polarization of F-actin and HLA-A, -B, -C is conserved in another breast cancer cell line. Emerald-LifeAct-expressing MDA-MB-468 cells (green) were pre-labelled for HLA-A,-B,-C (red) and conjugated with CTV-stained primary NK cells **(A-C)** or NK92-MI cells (cyan; **D-F)**. **(A and D)** Representative imaging flow cytometry images of cell-to-cell conjugates between NK cells and MDA-MB-468 cells with or without synaptic actin cytoskeleton remodelling at the immunological synapse, (AR+ and AR-, respectively). Scale bar: 10 µm. **(B and E)** Emerald-LifeAct relative intensity at the IS was used to classify MDA-MB-468 cells into AR+ and AR-groups (ratio >1 and ratio <1 respectively). Relative HLA-A, -B, -C intensities at the synapse in these 2 subgroups are presented. **(C and F)** The overall mean fluorescence intensity of HLA-A, -B, -C across the entire cell membrane of the cancer cell was measured in cell-to-cell conjugates formed between NK cells and AR+ or AR-MDA-MB-231 cells. **(B and C)** Data were collected from NK cells isolated from 1 donor, with n=100 cell-to-cell conjugates analysed per condition. **(E-F)** Data were collected from 1 experiment in each panel, with n=100 cell-to-cell conjugates analysed per condition. Statistical significance was determined using a Mann-Witney test.

**Supplementary Figure 4:**
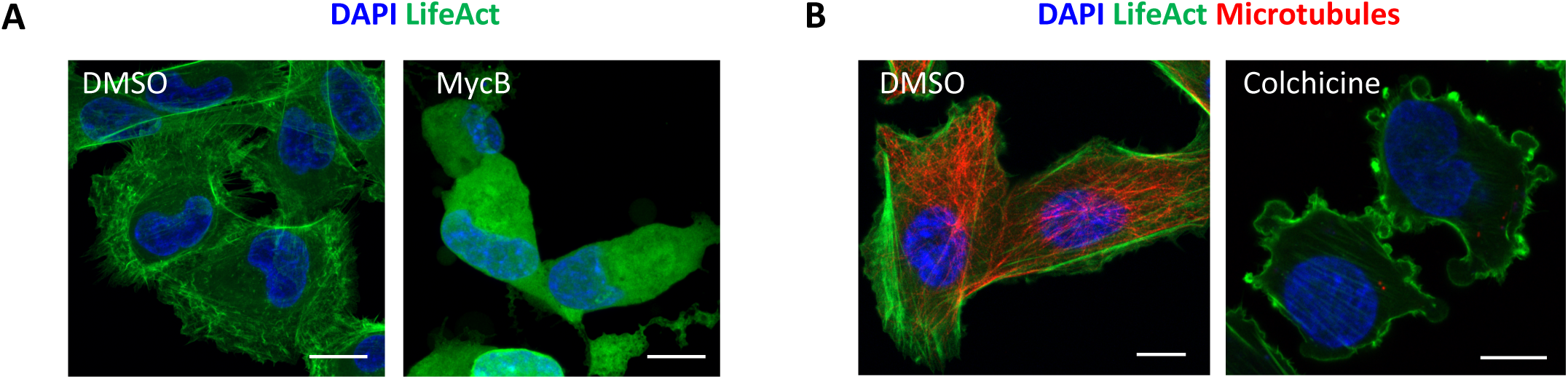
Control experiments for drug treatments affecting cytoskeletal organization in cancer cells. Adherent Emerald-LifeAct-expressing MDA-MB-231 cells (green) were treated with Mycalolide B (an inhibitor of actin filament polymerization) **(A)** or colchicine (a microtubule-disrupting agent) **(B)**. Following treatment, cells were washed, permeabilized and stained for the nucleus (blue) **(A and B)** and microtubules (red) **(B)**. Representative Airyscan images showing MDA-MB-231 cells with control (DMSO) or drug treatment. Maximum intensity projection images generated from 30 **(A)** or 20 **(B)** slices of a z-stack are shown. Scale bar: 10 µm.

**Supplementary Figure 5:**
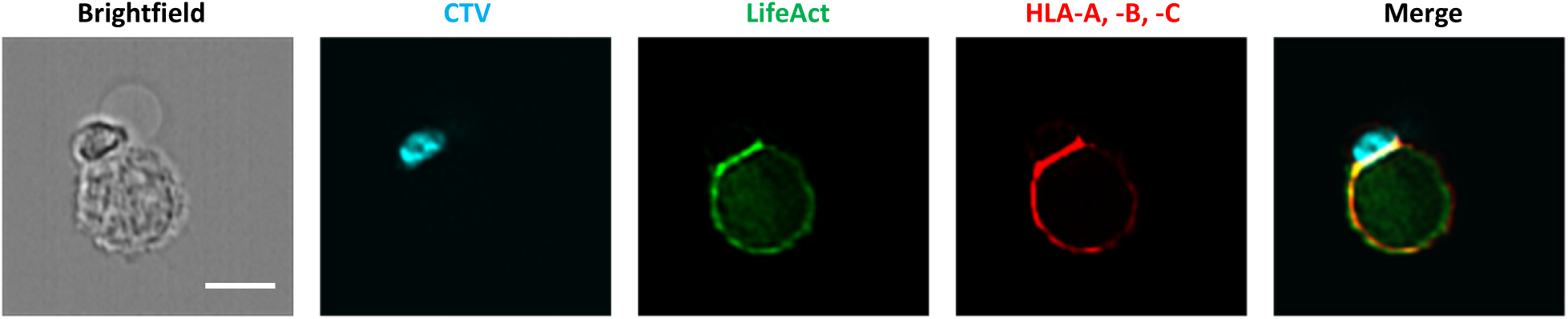
Control showing that HLA-blocking antibodies reveal a similar synaptic polarization pattern as non-blocking antibodies. Emerald-LifeAct-expressing MDA-MB-231 cells were pre-treated with either HLA-blocking antibodies or an isotype control antibody before being conjugated with CTV-labelled primary NK cells for 60 minutes. After conjugation, cells were incubated with a goat anti-mouse Alexa Fluor 555 secondary antibody (red). Scale bar: 10 µm.

**Supplementary Figure 6:**
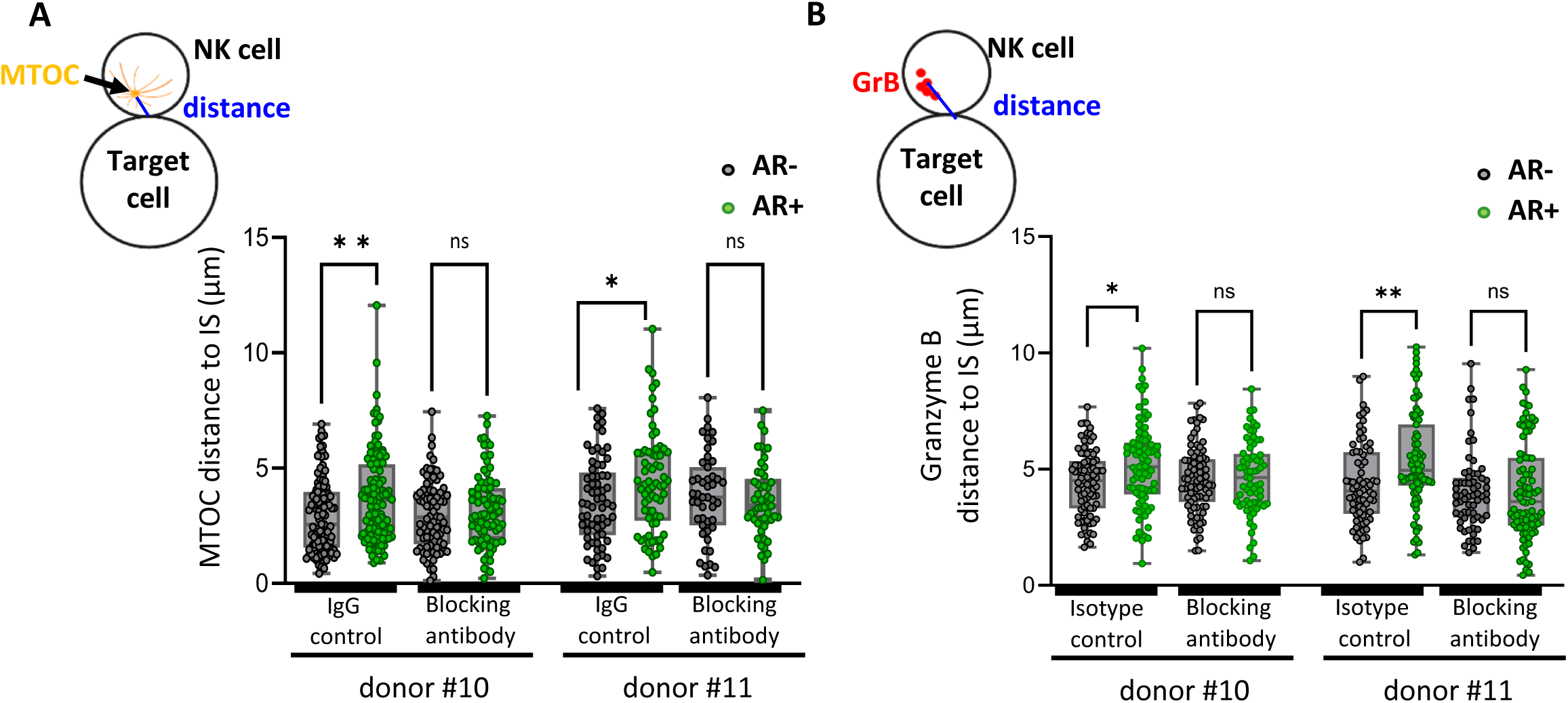
HLA-blocking antibody counteracts synaptic actin remodelling-driven inhibition of NK cell polarization in another breast cancer cell line. Emerald-LifeAct-expressing MDA-MB-468 cells were pre-treated with either HLA-blocking antibodies or an isotype control antibody before being conjugated with CTV-labelled primary NK cells for 60 minutes. After conjugation, cells were immunolabelled for granzyme B and γ-Tubulin and analysed using imaging flow cytometry. Emerald-LifeAct relative intensity at the immunological synapse was used to classify MDA-MB-468 cells into AR+ and AR-groups (ratio >1 and ratio <1 respectively). NK cell lytic machinery polarization was evaluated by measuring the distance between the MTOC and the IS centre **(B)**, as well as the distance between the Granzyme B centroid and the IS centre **(C)**. Distances for AR+ and AR-subgroups are presented. Data were collected from NK cells isolated from 2 distinct donors, with n=80 cell-to-cell conjugates analysed per condition. Statistical significance was determined using the Mann-Whitney test.

**Supplementary figure 7:**
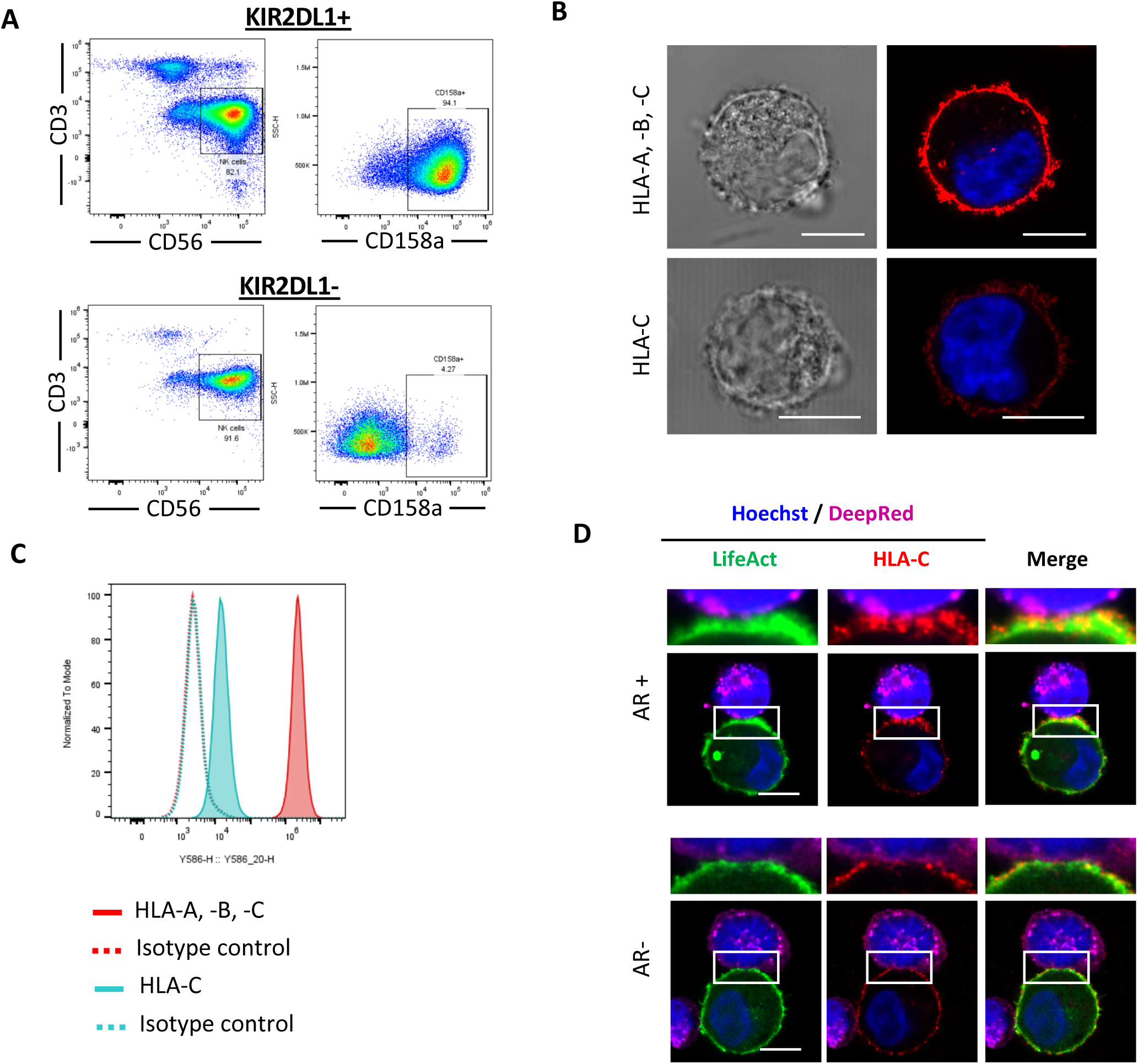
Charaterisation of KIR2DL1^+^ and KIR2DL1^-^ primary NK cells and HLA-C expression on MDA-MB-231 cells, conjugated or not with primary NK cells. **(A)** KIR2DL1 expression was analysed by flow cytometry in primary NK cells after sorting and amplification over two weeks prior to experimental use. The first panel shows CD56 and CD3 expression, while the second panel illustrates KIR2DL1 expression in CD56^+^ CD3^-^ cells. **(B)** MDA-MB-231 cells were pre-labelled with antibodies against HLA-A, -B, -C or HLA-C (red) and stained with Hoechst to visualize nuclei (blue), followed by confocal imaging. **(C)** Flow cytometry analysis of HLA-A, -B, -C and HLA-C expression on MDA-MB-231 cells. MDA-MB-231 cells were stained with PE-conjugated anti-HLA-A, -B, -C or anti-HLA-C antibodies and compared to the respective PE-conjugated isotype controls. Representative histograms display the fluorescence intensity of HLA-A, -B, -C and HLA-C relative to the isotype control. **(D)** Emerald-LifeAct-expressing MDA-MB-231 cells (green) were pre-labelled for HLA-C (red) and co-cultured for 40 minutes with Deep-Red-stained primary NK cells (purple) along with Hoechst staining for the nucleus (blue). Representative Airyscan images show cell-to-cell conjugates formed between primary NK cells and MDA-MB-231 cells with or without synaptic actin cytoskeleton remodelling (AR+ and AR-, respectively). Scale bars: 10 µm.

